# The dorsal hippocampus’ role in context-based timing in rodents

**DOI:** 10.1101/2022.01.10.475732

**Authors:** Benjamin J. De Corte, Sean J. Farley, Kelsey A. Heslin, Krystal L. Parker, John H. Freeman

## Abstract

To act proactively, we must predict when future events will occur. Individuals generate temporal predictions using cues that indicate an event will happen after a certain duration elapses. Neural models of timing focus on how the brain represents these cue-duration associations. However, these models often overlook the fact that situational factors frequently modulate temporal expectations. For example, in realistic environments, the intervals associated with different cues will often covary due to a common underlying cause. According to the ‘common cause hypothesis,’ observers anticipate this covariance such that, when one cue’s interval changes, temporal expectations for other cues shift in the same direction. Furthermore, as conditions will often differ across environments, the same cue can mean different things in different contexts. Therefore, updates to temporal expectations should be context-specific. Behavioral work supports these predictions, yet their underlying neural mechanisms are unclear. Here, we asked whether the dorsal hippocampus mediates context-based timing, given its broad role in context-conditioning. Specifically, we trained rats with either hippocampal or sham lesions that two cues predicted reward after either a short or long duration elapsed (e.g., tone-8s / light-16s). Then, we moved rats to a new context and extended the long-cue’s interval (e.g., light-32s). This caused rats to respond later to the short cue, despite never being trained to do so. Importantly, when returned to the initial training context, sham rats shifted back toward both cues’ original intervals. In contrast, lesion rats continued to respond at the long cue’s newer interval. Surprisingly, they still showed contextual modulation for the short cue, responding earlier like shams. These data suggest the hippocampus only mediates context-based timing if a cue is explicitly paired and/or rewarded across distinct contexts. Furthermore, as lesions did not impact timing measures at baseline or acquisiton for the long cue’s new interval, our data suggests that the hippocampus only modulates timing when context is relevant.

## Introduction

Learning and memory benefit individuals by allowing them to forecast future events. By drawing upon learned information, observers can adjust their behavior based on predictions over what will happen in the future. A core component of these predictions is anticipating *when* events will occur. Despite this, we have a limited understanding of how the brain regulates behavior based on time.

A variety of simple tasks can be used to study timing. They often involve presenting a subject with a cue that instructs them to make a motor response after they believe a certain interval of time has passed (Bader & Wiener, 2021; Balcı & Freestone, 2020). For example, investigators might present a rat with a tone that instructs them to press a lever after 8 seconds pass to earn reward. These tasks have provided valuable insight into how the brain mediates timing in both human (Coull et al., 2013; Damsma et al., 2021) and non-human animals (Matell et al., 2011; Mello et al., 2015; Wang et al., 2020). As a result, many neural models of timing center on explaining these data (Matell & Meck, 2004; Simen et al., 2011; Wang et al., 2018).

Given their simplicity, these tasks likely fail to capture how the timing system operates in real-world contexts. Of course, to study any cognitive process under controlled laboratory conditions, such task-simplifications will always be required. However, many attempts to adapt timing tasks to better capture the complexities of everyday life have led to the discovery of behavioral effects that conflict with existing neural models of timing (Balci et al., 2009; De Corte & Matell, 2016; Gür et al., 2018; Matell & De Corte, 2016; Matell & Henning, 2013; Raphan et al., 2019). For example, we recently proposed the ‘common cause hypothesis’—an evolutionary framework for conceptualizing how the timing system facilitates survival in realistic scenarios (De Corte et al., 2018). Its premise is best illustrated with the following analogy. Consider an animal that is foraging in an area that contains two plants—plant A and plant B. When searching through plant A, it typically finds food every 8s. In contrast, when searching through plant B, it finds food every 16s. These plants can be conceptualized as temporal cues, similar to ones that might be used in a laboratory experiment. Importantly, while these cues signal different durations, their intervals will often covary as environmental conditions change. For example, during a drought, both plants would produce less food than normal. Consequently, both of their durations will increase (i.e., food would be found less frequently/at longer intervals than normal).

Now, consider what would happen if the animal entered the drought-impacted area and learned that plant A’s interval increased. If the animal anticipates that plant A and B’s durations will covary, it can infer that plant B’s interval also increased, without physically visiting the plant itself. From an evolutionary standpoint, this would be highly advantageous because the animal could begin estimating the rate of return for foraging all plants in the area based on a change in one interval. Therefore, it could decide to stay or leave the location faster, relative to learning each plant’s new duration individually.

The essential point illustrated above is that, in naturalistic contexts, the intervals associated with different temporal cues will often covary due to a common underlying cause (e.g., weather conditions, seasonal changes, etc.). According to the common cause hypothesis, observers capitalize upon this covariance, such that, when one cue’s interval changes, temporal expectations for other cues shift in the same direction. Recently, we found evidence of this mechanism in a series of experiments (De Corte et al., 2018). As a basic test, we trained rats to associate two cues with different delays to reward. Then, we changed the interval associated with one cue (e.g., light-32s), and asked whether this would impact responding to the other cue. As predicted, rats responded as if they expected the other cue’s interval had changed in the same direction.

Importantly, seeing learning ‘transfer’ across cues conflicts with existing models of timing, as they typically assume different cues are processed independently (for discussion see De Corte et al., 2018). Therefore, it is important to begin establishing the neural mechanisms of this effect—our goal here. Another experiment provides a clear starting point. To illustrate the hypothesis, note that conditions will often differ between environments. A drought in North America does not typically tell us about the climate in Europe. Therefore, if a cue’s interval changes in one environment, observers should not necessarily assume that its interval has changed in other contexts. From this, we predicted that learning for a cue, as well as transfer to other cues, would be context-specific (De Corte et al., 2018, see Experiment 5).

To test this, we trained rats to associate two cues with different intervals in one context (e.g., tone-8s / light-16s). Then, we transitioned the rats to a new context and extended the interval associated with one of the cues (e.g., light-32s). As predicted, rats responded later to both cues when tested in this context. More importantly, when tested in the original training context, they shifted back toward the cues’ original intervals, indicating context-specificity.

Notably, the dorsal hippocampus is widely implicated in allowing context to modulate conditioned behavior in variety of non-timing tasks (Frankland et al., 1998; Penick & Solomom, 1991; Wiltgen et al., 2006). Here, we asked whether dorsal hippocampus lesions would prevent context-based timing in the above design. While this was our primary goal, we were also able to explore the hippocampus’ role in baseline timing performance and learning new intervals, which is also unclear. As such, we were able to assess the hippocampus in relation to the common cause hypothesis specifically, as well as its general role in timing.

## Methods

### Apparatus

We used custom operant chambers described in detail previously (Broschard et al., 2021). Briefly, the chambers were 35cm X 41cm X 36cm in width, depth, and height, respectively. The front wall contained a rectangular opening, where we mounted a 38cm LED monitor (Hewlitt-Packard) such that only the screen was visible from within the chamber. We also mounted a custom nosepoke below the bottom edge of the screen. The nosepoke consisted of a 3D-printed rectangular case (shielded with aluminum) with a 28mm hole through the center. The sides of the hole contained an infrared detector (Adafruit, 5V) oriented parallel to the floor, for detecting snout-entry. We used a custom feeder to dispense 45mg food pellets (Bioserv) into a food magazine located along the back wall. A tone generator (Sonalert, 4kHz, 80dB) and houselight (28V, 11 lux) served as cues during the experiment. To mask external noise, we placed each chamber in a sound-attenuating box with front panel-doors. We lined the internal walls and doors of the case with black sound proofing material (mass-loaded vinyl). Furthermore, we played white or brown noise continuously at 60dB (see below) through computer speakers mounted at opposing walls inside of the box. We accomplished stimulus control and data acquisition using custom Matlab scripts in addition to the Matlab-Arduino extension package. For stimulus/feeder control, we sent digital outputs to the Arduino, which triggered the desired component via a solid-state relay board (Sainsmart, 8-channel). The infrared detector in the nosepoke served as a digital input signal for detecting response onset/offset.

### Behavioral training

All protocols followed from those used in De Corte et al. (2018) as closely as possible. We trained male Sprague Dawley rats (n = 26) under modest food deprivation, matching the strain, sex-makeup, and deprivation conditions of our initial work. We monitored and maintained body weights at 85% of non-deprived levels. Initial training progressed through three phases. During the initial ‘shaping’ phase (3 sessions), rats learned to make responses on the nosepoke to earn food reward. Sessions lasted 60 minutes or until rats earned 60 rewards. We did not introduce cues during this phase, and any response on the nosepoke yielded reward. Next, a ‘fixed-interval’ training phase began (7 sessions; 120 mins each). During this phase, each trial began with the presentation of either the tone or houselight. Each cue predicted reward availability after either an 8 or 16 second duration elapsed. We counterbalanced the modality-duration relationship (e.g., tone-8s / light-16s or light-8s / tone-16s). As soon as the rat made a response after the respective cue’s interval passed, the cue terminated and reward was delivered. We did not penalize or reward early responses. All trials were followed by a dark intertrial interval (ITI), lasting 90s on average (30s minimum + a variable, 60s geometrically distributed interval). After fixed-interval training, a ‘probe training’ phase began (13 sessions; 120 mins each). Probe training was identical to fixed-interval training. However, during a subset of ‘probe trials’, the cue remained on for 96-128s, regardless of when rats responded, and no reward was delivered. Probe trials were randomly intermixed with rewarded trials. We gradually increased the proportion of probe trials during this phase (30% for the first 3 sessions and 50%, thereafter). Furthermore, we gradually decreased the ITI during this phase (30s-fixed + 30s-geometric for the first two sessions and 30sfixed + 15s-geometric, thereafter). One sham rat failed to acquire temporally controlled responding by the end of this phase even with extended training and was, therefore, excluded from the experiment.

Once trained, rats began a ‘change phase’, in which we changed the long cue’s interval from 16s to 32s. We did not present the short cue during this stage. Importantly, rats completed the training and change phases in separate contexts. For example, the two phases took place in separate operant chambers, located in the same training room. Furthermore, the two contexts differed with respect to several sensory features. First, during the training phase, we presented alternating black and white stripes (1cm) on the monitor mounted to the front wall. During the change phase, we presented a blank, grey screen instead, with the pixel value corresponding to the mean of the stripe image. Second, during the training phase, the chamber floor was made of a grid of stainless-steel bars (1cm X 1cm). In contrast, during the change phase, we covered the grid with sheet-metal drilled with evenly-spaced, circular holes (1cm diameter). Third, during the training phase, we dispensed vanilla-extract (McCormick, 0.4ml) on a Kim-Wipe and introduced it into the chamber by sliding it underneath the floor-grid, resting it under the food port. During the change phase, we replaced the scent with peppermint oil (Nature’s Truth, 0.4ml). Finally, during the training phase, we played continuous white-noise outside of the chamber, and during the change phase, we switched to brown-noise. All rats began to show stable peak-shaped responding near the new interval within 16 sessions, on average, with only minor deviations (+/- 3 sessions).

Finally, after rats adapted to the new interval, the test phase began where we evaluated how rats would respond to either cue in the original training context, compared to the change-context. We conducted one test session in each context, including 12 probe trials for each cue (randomly intermixed) during each session. We also counterbalanced the order of testing in each context across rats (i.e., train-context vs. change-context first).

### Surgery and Histology

One week before the start of training, we administered rats with either lesions to the dorsal hippocampus (*n* = 13) or sham lesions (*n* = 12) to this area. Specifically, we anesthetized rats with isoflurane, using 5.0% for induction and 2.5% for maintenance. Then, we retracted the scalp and lowered a 30-gauge infusion cannula (Component Supply, Inc) at four sites in the dorsal hippocampus, spanning its anterior and posterior extent (Anterior, AP: -3.0, ML: +/-1.8, DV: -3.7; Posterior, AP: -4.2; ML: +/-3.2; DV: -3.9; relative to bregma at skull surface). We connected the cannula to an infusion pump (World Precision Instruments) with polyethylene tubing. After lowering the cannula to each site, we infused .3ul of NMDA (20 mg/ml; Millipore-Sigma) or saline, at 0.1μl/min. Following each infusion, we left the cannula in place for five minutes to allow for diffusion. After the experiment, we perfused rats using chilled saline followed by 4.0% paraformaldehyde (120ml each). Brains were extracted, post-fixed in paraformaldehyde (48hrs), equilibrated in 30% sucrose, sliced at 40μm sections, stained with thionin, and then imaged under a brightfield microscope to verify lesion locations.

### Behavioral analysis

We analyzed mean response rates by grouping responses during probe trials into 1s bins and averaging across trials. We fit each response curve with a Gaussian + kurtosis parameter function described in the following equation (De Corte et al., 2018; Swanton et al., 2009):

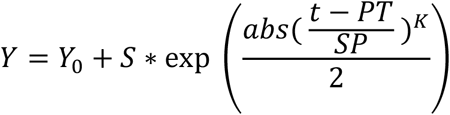

*Y*_*0*_ represents the baseline rate of responding, *S* is a scale parameter, *t* is time, *PT* is the mean (i.e., ‘peak-time’), *SP* is the spread, and *K* allows the function to accommodate varying degrees of skew. We extracted the peak-time, coefficient of variation (CV; spread / mean), response rate (i.e., peak-amplitude), and fit-quality (*R*^2^) to quantify performance. For *R*^2^ analyses, we used the Fisher’s *R* to *Z* conversion on the square root of each value (i.e., the correlation coefficient). This is standard and required to obtain normality in data based on Pearson-correlations. To filter-out trials with non-temporally controlled responding, at least one response had to occur before 64 seconds had passed during the trial to be included (i.e., twice the longest duration used in the experiment of 32 seconds). This criteria is highly conservative, relative to other approaches used for this task (e.g., Church et al., 1994).

We also analyzed single-trial patterns of responding, again using 1s bins. Specifically, we identified bursts of responses using a standard approach in which a dual step-function (i.e., rectangular pulse: first step-up, second step-down) is fit to response rates across time during a given trial (Church et al., 1994; De Corte et al., 2021). The algorithm iteratively fits the step function across time, finding the points where absolute residuals are minimized. Once identified, the time of the first step-up and second step-down are taken as the start and stop time, respectively. In line with response-rate analysis, we excluded trials where the start time occurred after twice the peak-time had passed (Matell et al., 2016).

### Statistical analysis

In all cases, we used a mixed-model Analysis of Variance as an omnibus test on each measure. We always included Group (lesion vs. sham) and Modality-Duration Pairing (tone-8s / light-16s vs. light-8s / tone-16s) as between-subjects factors. For the baseline phase, Cue (8s vs. 16s) served as a within-subjects variable, and data came from the last training session. For the change-phase, we divided the data into 16 equally-spaced blocks—corresponding to the mean number of acquisition sessions—to factor out slight differences in the number of retraining sessions across rats (i.e., Vincintized the data; Broschard et al., 2019). We split the data into an ‘Early’ and ‘Late’ stage, corresponding to the first and last 8-blocks, respectively. We tested each stage separately, and Block served as the within-subjects factor. For the test phase, Cue and Context (training, change, original) served as within-subjects factors. For Context, training refers to the last training day, and change/original refer to data from the two test sessions. We used an alpha level of 0.05. We assessed sphericity with Mauchley’s test and, when violated, used the Greenhouse-Geisser correction. Where appropriate, we conducted planned comparisons with t-tests.

## Results

### Overview of behavior and design

Environmental conditions will often cause the durations associated with different temporal cues to covary. Under the common cause hypothesis, observers assume that, when one cue’s interval changes, the durations associated with other cues have changed as well, due to a common causal factor (De Corte et al., 2018). Importantly, as conditions will differ across distinct environments, changes in temporal expectations should be specific to contexts where a duration-change has occurred. We asked whether the context-sensitivity and/or transfer depends on the dorsal hippocampus.

Specifically, we trained rats with dorsal hippocampus lesions (*n* = 13; Fig 1 C) or sham lesions (*n* = 12; Fig 1 C) on a timing task called the ‘peak-interval procedure’ (Fig 1 A,B). During this task, trials begin with the presentation of a cue that instructs rats to respond after a specific time-interval has passed to earn reward (e.g., tone-8s). ‘Probe trials’ are intermixed where the cue remains present for much longer than normal and no reward is provided. During probe trials, rats’ responses cluster around the entrained interval, allowing us to measure the cue’s temporal expectation. We trained rats to associate two cues with distinct durations (e.g., tone-8s / light-16s). Once trained, we placed rats in a new context and changed the long cue’s interval from 16s to 32s. We did not present the short cue during this phase. This allowed us to test the prediction that increasing the long cue’s interval will naturally increase to the short cue’s temporal expectation. Importantly, we also evaluated whether changes in either cue’s temporal expectation would be more dramatic in the context where the long-interval changed. We tested this in a final phase by presenting both cues in either context, expecting contextual modulation in the sham group but not the lesion group.

**Figure 1.**
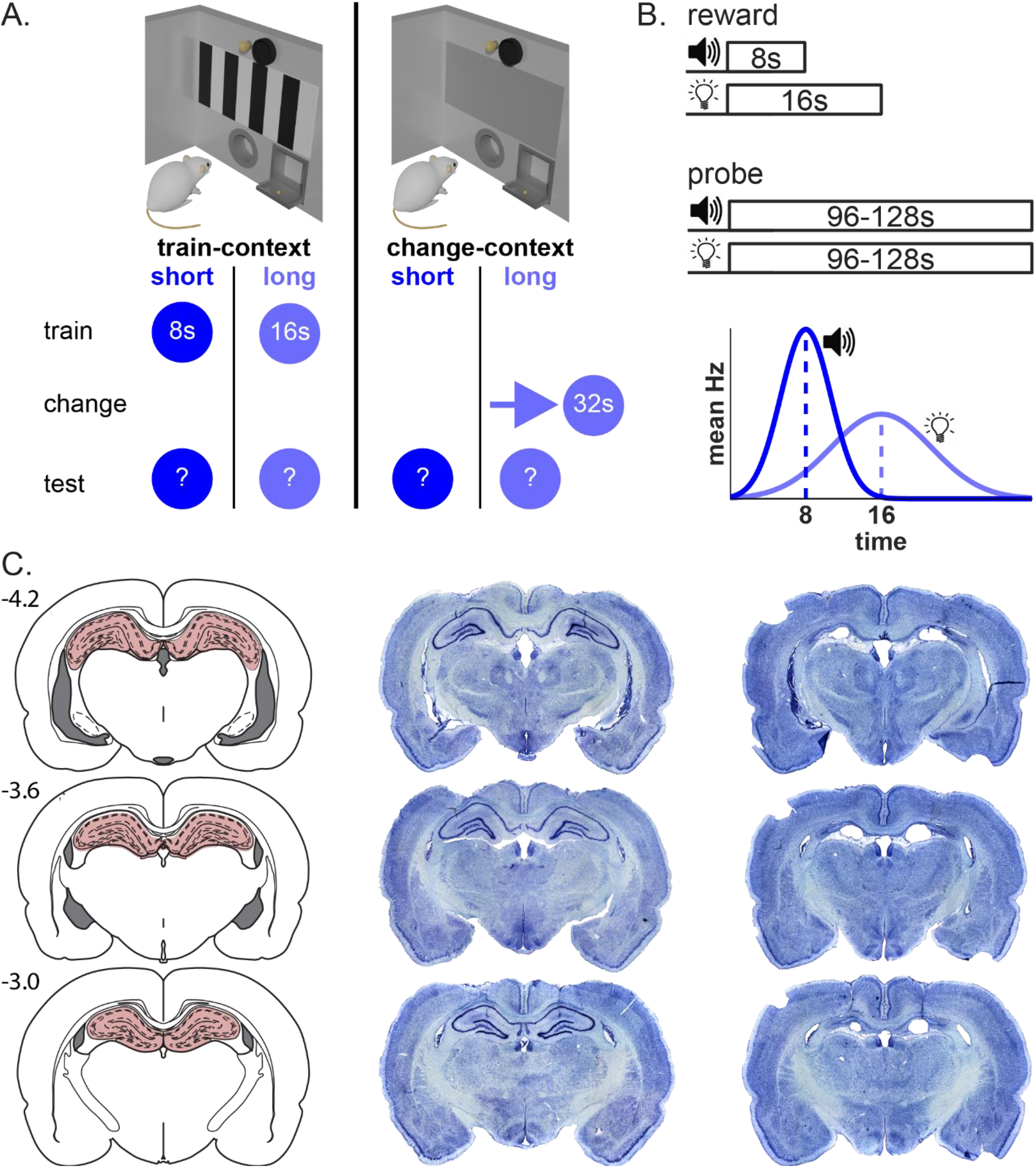
Task summary and histological illustration. (A) Task structure, including diagrams of the two contexts (top), and the intervals for the short and long cues across phases (bottom). Question marks during the test phase indicate that all trials were probes (i.e., no specific interval in effect) (B) Basic illustration of trial structure, focusing on the baseline phase. During rewarded trials, cue-termination/reward delivery occurs for the first response after the presented cue’s interval elapses. During probe trials, the cue is presented for an extended, variable amount of time and no reward is delivered. Bottom graph illustrates expected probe-trial behavior for each cue in trained rats. (C) Histology. Left column show atlas images at different anterior-posterior coordinates, with the dorsal hippocampus highlighted in red. Middle and right columns show a sham and lesioned rat, respectively.

### Effects of dorsal hippocampus lesions on baseline timing performance

First, we evaluated whether lesions impacted baseline timing during the training phase, in which rats associated the short cue with 8s and the long cue with 16s (Fig 2A). Figure 2B shows average probe trial responding for both cues during this phase. As expected, both groups showed gaussian-like response curves, with ‘peak times’ falling near each cue’s interval. Notably, no substantial group differences appeared to be present.

**Figure 2.**
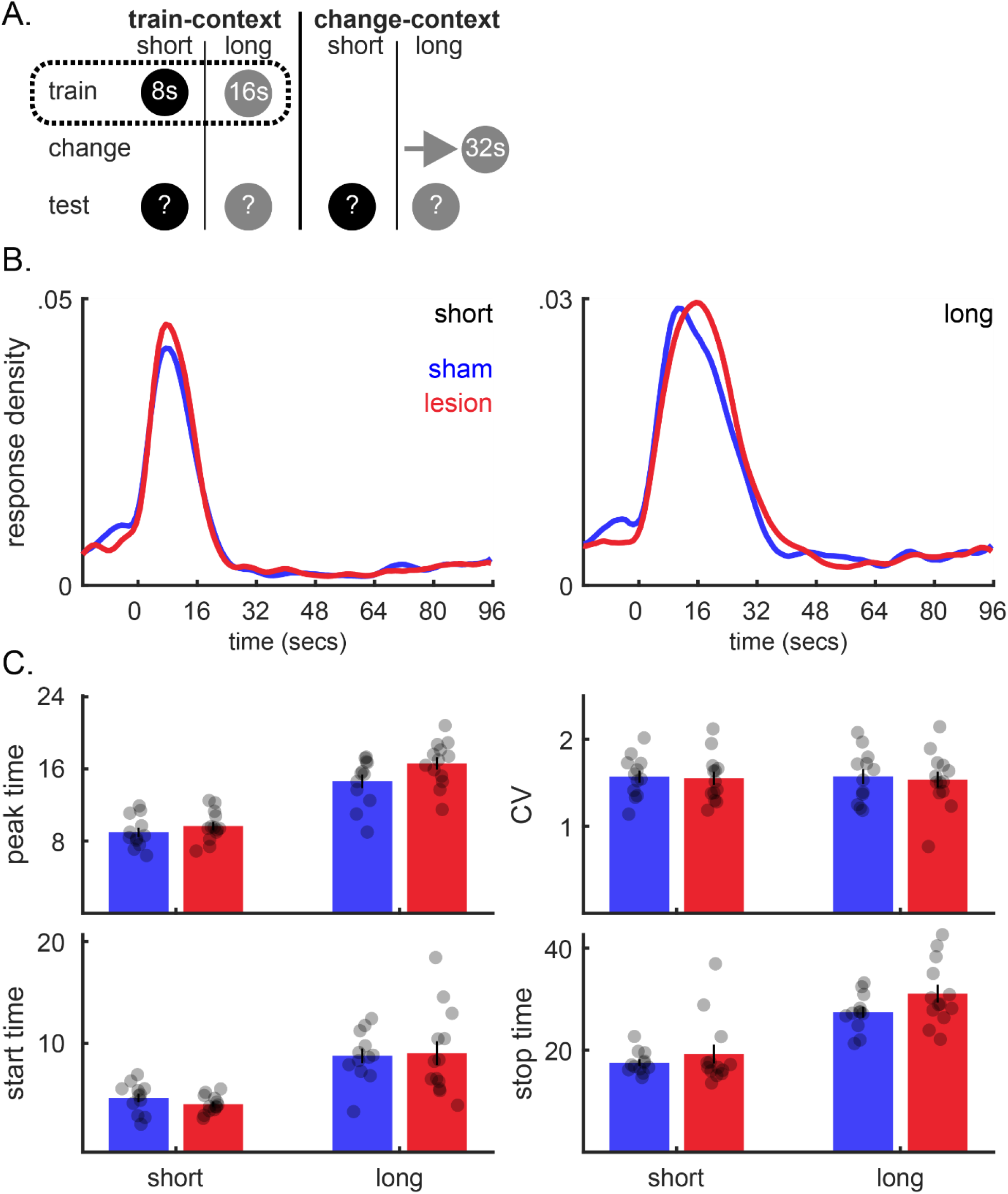
No notable differences in baseline timing performance between sham rats and rats with dorsal hippocampus lesions. (A) Minimal task diagram. Dotted rectangle emphasizes data come from the initial training phase. (B) Mean probe trial response rates under baseline conditions. Left and right panels show data from short- and long-cue trials, respectively. For presentation, data are normalized to the sum of responding and smoothed over a 3-bin window. (C) Measures of timing under baseline conditions. Peak-time and CV (spread / mean) are derived from fitting mean response distributions with Gaussian-like functions. Start and stop times are derived from analysis of single-trials.

To quantify this, we first fit each response distributions with a Gaussian function. Our primary measures of interest from the fits were the peak-time (mean) and CV (spread / mean)—measures of timing accuracy and variability, respectively. We did not find group differences for either measure [Fig 2C; Peak time: *F*s < 2.8, *p*s > 0.05; CV: *F*s < 1.0, *p*s > 0.05]. Furthermore, CVs remained relatively constant across intervals, indicating both groups showed the scalar property of timing, wherein the spread increases in constant proportion with the peak-time (Buhusi et al., 2018) [Fig 2C; Cue: *F*(1,21) = 0.02, *p* = 0.899]. We also did not find differences in overall fit-quality (i.e., *R*^2^), suggesting no substantial differences in the overall shape of responding were present during training [*F*s < 2.5, *p*s > .05]. Finally, lesions did not reliably affect motivation or motor function, as peakrates were comparable across groups [*F*s < 2.7, *p*s > 0.05].

Importantly, during individual trials in this task, rats do not respond in a smooth Gaussian manner, as suggested by the average response curves. Rather, they emit a sharp burst of responses that clusters around the target interval. Slight trial-to-trial variation in when the burst occurs leads averaged response rates to appear Gaussian. When the burst starts and stops provides further measures of timing performance (Buhusi et al., 2018).

Consistent with the mean-response measures, we did not find reliable group differences for start or stop times [Fig 2D; Start: *F*s < 1.5, *p*s > 0.05; Stop: *F*s < 3.3, *p*s > 0.05].

Collectively, these data suggest that the dorsal hippocampus is not critical to baseline timing performance during this task.

### Effects of dorsal hippocampus lesions on acquisiton in a new context

Next, we analyzed data from the change phase, in which we moved rats to a new context and changed the long cue’s duration from 16 to 32 seconds (Fig 3A). We did not present the short cue during this phase. Figure 3B shows mean response distributions at early, middle, and late periods of acquisiton for the two groups. Figure 3C and 3D shows how different performance measures changed across acquisiton blocks. For visual comparison, we also plot baseline data from the training phase (first value on each x-axis).

**Figure 3.**
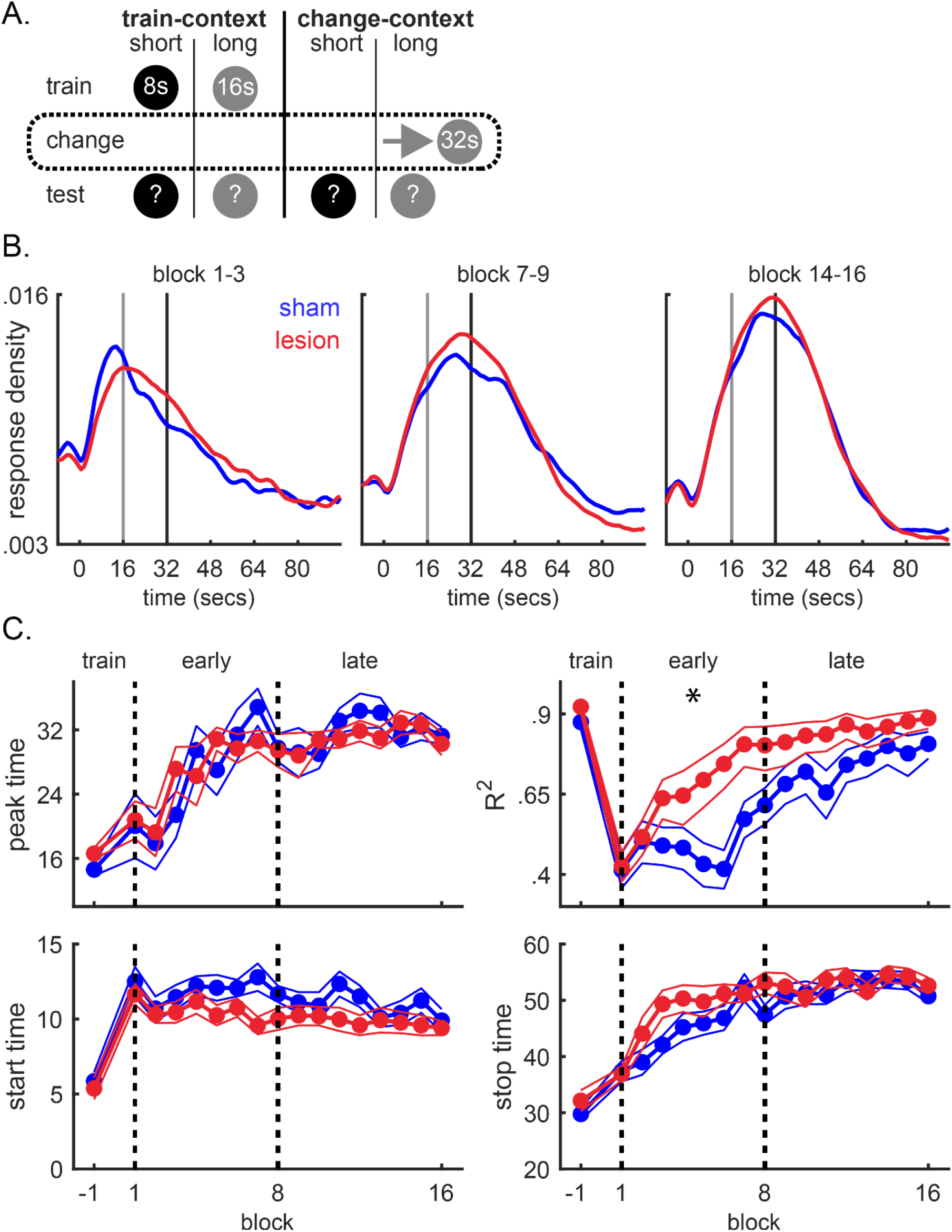
Effects of dorsal hippocampus lesions on acquisition of a new interval following a context change. (A) Minimal task diagram. (B) Average probe trial response rates at different points of acquisiton. Data are pooled across 3 -block windows occurring at the start, middle, and at the end of the acquisiton phase (left, middle, right, respectively). (C) Performance measures across individual acquisition blocks, including the peak-time, *R*^2^ of the Gaussian fit, start times, and stop times. For reference, the first value on each axis shows data from the last training session. Dotted lines demarcate phases of acquisition, defined as the ‘early’ and ‘late’ stages. Asterisks indicate significant group-differences during that stage. Other effects (e.g., block) are noted in-text.

Consistent with the baseline data, we did not find notable group differences for measures of timing accuracy. For example, peak-times adjusted rapidly, effectively stabilizing by the late-stage of acquisition [Fig 3C; Early: Block, *F*(7,147) = 9.60, *p* < 0.001; Late: Block, *F*(7,147) = 1.48, *p* = 0.215]. Importantly, we did not observe significant effects or interactions between groups [*F*s < 1.10, *p*s > 0.05]. Start and stop times followed the same pattern. Specifically, start-times stabilized quickly, appearing to level-off during the early phase [Fig 3D; Early: Block, *F*(7,147) = 1.33, *p* = 0.239; Late: Block, *F*(7,147) = 1.74, *p* = 0.105]. Similar to peak-times, stop times gradually adjusted during the early stage and stabilized by the late-stage [Fig 3D; Early: Block, *F*(7,147) = 18.03, *p* < 0.001; Late: Block, *F*(7,147) = 1.71, *p* = 0.111]. Importantly, we did not find group differences for either measure across acquisition [*F*s < 3.48, *p*s > 0.05].

While timing-accuracy adjusted comparably across groups, we did begin observing signs of group-differences during this phase of the experiment. Specifically, early in acquisiton, response distributions for both groups showed a positively skewed shape that extended broadly across the trial period (Fig 3B, left panel). Consistent with this, fit-quality of the Gaussian dropped in both groups early in acquisiton and ascended by the late stage [Fig 3C; Early: Block, *F*(7,147) = 9.05, *p* < 0.001; Late: Block, *F*(7,147) = 5.74, *p* < 0.001].

Surprisingly, the shape of peak-responding recovered faster in the hippocampal lesion group, particularly during the early-stage [Early: Block X Group, *F*(7,147) = 3.17, *p* < 0.05; Late: Block X Group, *F*(7,147) = 0.617, *p* = 0.742].

This disruption could reflect adjustments in timing toward the new interval and/or a general disorganization in behavior from the context-change itself (e.g., noise from increased exploratory responses). Timing adjustments toward the new interval should increase the variability of our timing measures (i.e., CV), with the effect reducing faster in the lesion group. As one might expect, the breadth and skew of the distributions led to volatile CV estimates from the fits (Fig S1). Therefore, we supplemented this measure by analyzing the CVs of start and stop times, which proved less outlier-prone given that they segregate timed responses on a trial-by-trial basis. Both groups adjusted comparably across measures [Fig S1; Session X Group *F*s < 1.5, *p*s > 0.05]; with the only reliable effect being a decrease in stop CVs across the early block, regardless of group [Fig S1; Session: *F*(7,147) = 11.03, *p* < 0.005]. Together with the comparable timing-accuracy measures across groups, the data overall favor the interpretation that sham rats were more sensitive to the context-change, rather than showed a timing deficit, per se.

In relation to the baseline data, these results further indicate that hippocampal lesions do not impact timing accuracy or variability, even when adapting to new intervals. However, lesions might make rats more resistant to the effects of context-changes on the overall organization of behavior during this task, which parallels work in Pavlovian appetitive conditioning (see Discussion; Penick & Solomom, 1991).

### Effects of dorsal hippocampus lesions on contextual-modulation of timed responding

Finally, we analyzed our primary data of interest—how rats would respond to both cues when tested in either the original training context or the context in which the long cue’s interval changed from 16 to 32 seconds (Fig 4A). For clarity, we refer to these as the original context and change context, respectively. To avoid giving rats feedback regarding when to respond, we only included probe trials during this phase. For the sham group, we expected that shifts in timing for both cues would be more pronounced in the change-context, relative to the original context. In contrast, we did not expect to see contextual modulation in the hippocampal lesion group.

**Figure 4.**
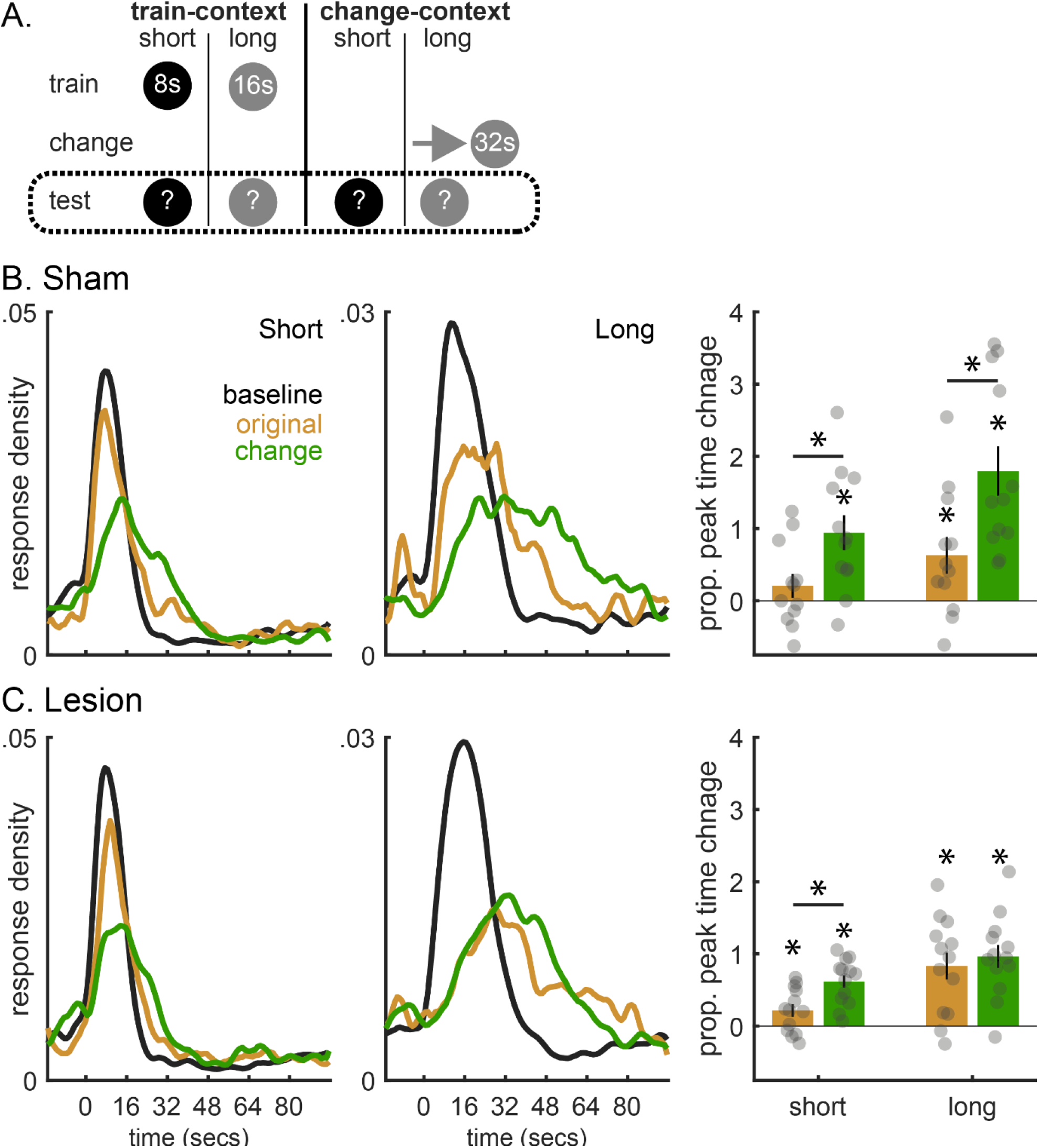
Dorsal hippocampus lesions impact context-based timing only for the explicitly changed-cue. (A) Minimal task diagram. Dotted rectangle emphasizes the primary data of interest come from the test phase. (B) Test data for the sham group. Average probe trial response rates for the short cue (left) and long cue (middle) during training, testing in the original context, and testing in the change context. The right panel shows proportional shifts in peak times during testing in either context, relative to baseline training. (C) Same as B, yet showing data from the lesion group. In all cases, asterisks centered over the bars indicate significant peak time shifts from baseline training, and asterisks spanning pairs of bars indicate significant differences between testing in either context.

All data from the sham group matched our predictions, as depicted in Figure 4B. For clarity, peak times are expressed as proportional shifts from initial training, with higher values indicating rightward shifts (Fig 4B, right panel). For the long cue, sham rats showed rightward shifts during testing in both contexts, suggesting some context-generalization did occur from the change-context to the original context [Fig 4B; Context: *F*(2,42) = 47.15, *p* < 0.001; train vs. original: *t*(11) = 2.65, *p* < 0.05; train vs. change: *t*(11) = 6.86, *p*

< 0.001]. Importantly, the shifts were more pronounced in the change-context, relative to testing in the original context [Fig 4B; *t*(11) = 6.23, *p* < 0.001]. This supports the common cause hypothesis’ prediction that shifts in temporal expectations should be modulated by the physical context in which a duration-change occurs. Critically, short cue responding paralleled these effects, being rightward shifted in the change-context, relative to both initial training and testing in the original context [Fig 4B; train vs. change: *t*(11) = 4.38, *p* < 0.005; original vs. change: *t*(11) = 2.58, *p* < 0.05]. The minor difference was that testing in the original context was not reliably different from training [*t*(11) = 1.34, *p* = 0.21]. As we always rewarded the short cue at 8s, these data support the common cause hypothesis’ prediction that, when one cue’s interval changes, the temporal expectations for other cues will shift in the same direction in a context-dependent manner.

The lesion group’s data are depicted in Figure 4C. For the long cue, rats showed rightward shifts in both contexts during testing, similar to the sham group [Fig 4C; train vs. original: *t*(12) = 4.50, *p* < 0.001; train vs. change: *t*(12) = 7.07, *p* < 0.001]. Importantly, unlike the sham group, these shifts did not vary reliably across contexts, consistent with our prediction [Fig 4C; Context X Lesion: *F*(2,42) = 5.31, *p* < 0.01; original vs. change: *t*(12) = 0.34, *p* = 0.737]. However, we were surprised to see contextual modulation for the short cue [Fig 4C; Cue X Context X Lesion: *F*(2,42) = 4.45, *p* < 0.05]. In line with the sham group, these shifts were greater during testing in the change-context, relative to both training and testing in the original context [Fig 4C; train vs. change: *t*(12) = 7.13, *p* < 0.001; original vs. change: *t*(12) = 2.99, *p* < .05]. The primary difference from the sham group was that testing in the original context produced reliable, rightward shifts [*t*(12) = 2.65, *p* < 0.05]. As this comparison was not significant for the sham group, this gives some indication of decreased contextual modulation for the short-cue in the lesion group. However, when we computed the difference between each subject’s proportional peak-time shifts during testing in either context for the short cue, we found no reliable difference between groups [*t*(23) = 1.10, *p* = 0.284]. In contrast, the long-cue differences were significantly greater in shams [*t*(23) = 3.28, *p* < 0.005]. Overall, these data support our prediction that the hippocampus regulates context-based timing, although the degree of its involvement unexpectedly differed across cues.

Unfortunately, extinction occurs rapidly during this task and increases noise in any form of operant or Pavlovian behavior (De Corte & Matell, 2016; Gharib et al., 2001; Stahlman et al., 2010). As testing was conducted in extinction (i.e., all probes), we were unable to obtain robust estimates of single-trial behavior (i.e., start/stop times), similar to our prior work (De Corte et al., 2018). However, in the supplemental results section, we provide an exploratory single-trial analysis, and note that the primary findings are generally consistent with the peak-time data. However, they might suggest attenuated contextual modulation for short cue start times in the lesion group (Fig S2).

As a whole, these data suggest that, while the dorsal hippocampus plays a negligible role in timing at baseline or during acquisition, it will begin modulating the timing network when context becomes relevant. However, given the presence of contextual modulation for the short cue at test and lack thereof for the long cue in the lesion group, this area may play a more complex role in context-conditioning than generally believed.

## Discussion

The common cause hypothesis predicts that, when the interval associated with one cue changes, temporal expectations for other cues will shift in the same direction (De Corte et al., 2018). Importantly, updates to any temporal expectation should be more prominent in contexts in which a duration-change has occurred. We asked whether the dorsal hippocampus mediates this context effect.

First, we trained rats with either dorsal hippocampus lesions or shams to associate two cues with either an 8s or 16s duration. Then, we transitioned rats to a new context and increased the long-cue’s interval to 32s. Finally, we tested how rats would respond to either cue in the initial training context or the context in which the long-interval changed. Relative to training, sham rats responded later to both cues in the context where the long cue’s interval changed. Critically, these shifts were attenuated during testing in the initial context. This replicates our prior behavioral data, providing further support for the common cause hypothesis (De Corte et al., 2018; Matell & Henning, 2013). Importantly, consistent with our prediction, lesioned rats did not show reliable contextual modulation for the long cue, always persisting at its new interval. However, they did show contextual modulation for the short cue, responding earlier in the initial training context like shams. These data contribute to our understanding of the hippocampus’ role in timing and conditioning more broadly.

For example, lesions blocking contextual modulation for the long cue gives first-insight into the neural mechanisms underlying the common cause hypothesis—our primary goal. Most neural theories of timing typically focus on explaining baseline timing performance. Consequently, they often have difficulty accounting for more complex timing phenomena, such as transfer or context-based timing (for discussion see: De Corte et al., 2018; Matell & Henning, 2013). Determining how to adapt existing models to account for the hippocampus’ role in this effect will require more sophisticated techniques (e.g., electrophysiology). However, the design-validation and causal data provided here constitute an important first-step toward this goal.

Moreover, the fact that, on a within-subject basis, lesioned rats showed reliable contextual modulation for the short-cue but not the long-cue opens several questions, both for timing and general context-conditioning. Presumably, the discrepancy relates to how the two cues differed with respect to the task-structure. For example, the first time the short cue was presented in the new context was during testing. In contrast, we presented the long cue repeatedly in both contexts during prior phases. Therefore, one possibility is that, for the hippocampus to impact contextual modulation, a given cue must be directly experienced in different contexts. To some degree, this matches with the hippocampus’ theorized role in memory. Specifically, theories of memory often pose that, when an observer experiences an event, fragments of the episode (e.g., sights, sounds) are stored in disparate cortical areas (Teyler & DiScenna, 1986; Teyler & Rudy, 2007). The hippocampus tracks where these fragments are stored and, at retrieval, reactivates them, binding the components into a unified representation (Jadhav et al., 2016; M. A. Wilson & McNaughton, 1994). In our task, binding would be possible for the long cue (i.e., long cue + new context vs. long cue + old context). However, as the short cue was never paired with the new context before testing, this would not be possible. This might encourage hippocampal recruitment for the long cue, with other areas engaging for the short cue.

This interpretation agrees with work suggesting that the hippocampus is selectively recruited to reduce interference between related memories that must be distinguished based on context (Agster et al., 2002; Butterly et al., 2012; Smith & Bulkin, 2014). For example, Bulkin et al., 2016 recently trained rats on an odor-discrimination task where the task-contingencies changed mid-way through the study (e.g., choose scent A during initial sessions; avoid scent A during following sessions). When the contingency-change occurred in the same context as training, hippocampal cells fired similarly before and after the update, and rats showed proactive interference. In contrast, when the reversal occurred in a novel context, hippocampal units (both spatial- and task-sensitive) remapped their activity, which appeared to reduce proactive interference. Prior causal work with this task had already shown that the hippocampus is only necessary for performance when context-changes occur (i.e., the latter group; Butterly et al., 2012). Therefore, Bulkin et al., 2016 concluded that, when the same cue (e.g., scent A) must be associated with different memories in two contexts, the hippocampus selectively engages to reduce interference between the two memories across environments. Related to our task, pairing the long-cue in the two contexts would encourage hippocampal interference-reduction, which would not be possible for the short-cue where information transfers via a more ‘inferential-like’ process because rats were never trained to shift their responses.

A related difference is that we explicitly rewarded the long cue at its new interval in the change-context and at its original interval in the original-context. Consequently, when testing began, rats had received unambiguous instruction regarding when to respond to the long cue in both contexts. In contrast, we provided more ambiguous information for the short cue; only rewarding it at its original interval in the training context. This opens the possibility that hippocampus’ involvement in context-effects varies as a function of a cue’s ambiguity. Interestingly, the same proposal has been made previously in an independent line of work. The data relate to a finding in Pavlovian conditioning experiments referred to as the ‘renewal effect’ (Bouton, 2004; Bouton et al., 2011).

In a simple renewal design, rats are initially trained that a cue predicts reinforcement in one context (e.g., context A: tone + shock). Then, they are placed in a new context and the CS-US association is extinguished (e.g., context B: tone + no shock). As expected, conditioned responses to the cue decrease in the new context. However, if rats are returned to the initial context, conditioned responses to the cue re-emerge (i.e., ‘renew’). This effect has been important for establishing that extinction does not ‘erase’ previously learned CS-US associations. Rather, extinction learning modulates the expression of behavior for a given cue, in a context-dependent manner.

Many have asked if the hippocampus mediates the renewal effect (Corcoran & Maren, 2001, 2004; Frohardt et al., 2000; A. Wilson et al., 1995). Similar to our case, the results depend on the task-structure. Specifically, different variants of the renewal design above have been developed, and hippocampal involvement depends on what variant is used. For example, as described above, rats can be trained in one context, extinguished in another, and tested in the initial training context—referred to as an ABA design. In this case, hippocampal manipulations have no effect on renewal (Corcoran & Maren, 2004; Frohardt et al., 2000). However, rats can also be trained in one context, extinguished in another, and then tested in an entirely novel context—called an ABC design. In this case, hippocampal manipulations block renewal, with responses being suppressed during testing (Corcoran & Maren, 2004).

To explain this difference, Corcoran & Maren (2004) argue that ambiguity mediates hippocampal involvement between task-variants. Specifically, in an ABA design, rats receive feedback about the cue in each context, keeping ambiguity low during testing. Conversely, in the ABC design, rats are never given feedback about the cue in the novel test context, providing higher ambiguity. As lesions only affect performance during ABC designs, Corcoran & Maren (2004) proposed that the hippocampus only mediates context effects when ambiguity is high.

Interestingly, our data favor the reverse possibility. We rewarded the long cue in both contexts, yielding low ambiguity, and hippocampal lesions impacted performance. Conversely, the short cue was more ambiguous, yet lesions did not affect context-based responding. Despite this, our data still suggest ambiguity (per se) affects hippocampal recruitment in context conditioning. Furthermore, our data add that task-contingencies might moderate this relationship (e.g., Pavlovian vs. operant, extinction vs. timing, etc.). Moreover, unlike renewal designs, our task incorporates two cues that differ in ambiguity, allowing for a within-subject exploration of this topic. One interesting experiment would be to run the equivalent of an ABC renewal design in our task, testing the rats in an entirely novel context. Predictions from the common cause hypothesis are less clear here. However, with high ambiguity for both cues, any context-modulation should not be hippocampal-dependent, contrasting with the renewal data.

Notably, both our data and the renewal effects suggest that other regions process context independently of the hippocampus. While we can only speculate as to what these areas might be, prior work suggests that cortical areas represent contextual variables, such as the human PFC (Waskom & Wagner, 2017) and the rodent medial frontal cortex (Hyman et al., 2012; Sharpe & Killcross, 2015; Zelikowsky et al., 2014). To our knowledge, the extent to which this contextual processing is contained within the cortical areas themselves is unclear. However, pursuing this question might suggest new targets regarding how temporal learning transfers across cues and/or its contextual modulation, which our data suggest are not fully hippocampal-dependent processes.

Our data also have important implications regarding whether the hippocampus mediates baseline timing. For example, the hippocampus is undoubtedly involved in encoding time in episodic memory (Tulving, 1986), and CA1 neurons appear to encode time during a variety of tasks (i.e., ‘time-cells’; Eichenbaum, 2014; Kraus et al., 2013). Naturally, many question whether this temporal coding contributes to timing during tasks like the one used here. Some work with the peak-procedure suggests that hippocampal lesions produce a persistent leftward shift in responding during baseline performance (Meck et al., 1984; Yin & Meck, 2014). For this reason, we used pre-training lesions, to keep learning/performance processes as constant as possible throughout the experiment. However, converging evidence suggests that the hippocampus is only recruited at baseline when exceptionally long intervals are used—on the order of several minutes. Specifically, when rodents time multi-second intervals, such as the ones used here, hippocampal effects are often absent or minor (Dietrich et al., 1997; Dietrich & Allen, 1998; Gupta et al., 2019; Jacobs et al., 2013). However, when rodents track intervals lasting several minutes, hippocampal manipulations begin to exert an effect (Jacobs et al., 2013; Shikano et al., 2021). Similarly, patient H.M., who received bilateral hippocampus resections, accurately timed multi-second intervals (e.g., ∼20 seconds), yet began underestimating intervals in the minutes-range (e.g., ∼5 minutes; Richards, 1973).

Our data add to this literature, as we did not observe baseline effects of lesions. Furthermore, during the change-phase, lesioned rats adapted normally to the long cue’s new interval with respect to all measures of timing accuracy and variability. However, early in acquisition, we did find a general disruption in the overall shape of peak responding that recovered more quickly in lesioned animals. Given the comparable timing measures across groups, the most plausible explanation is that the context-change itself produced a general disruption to operant behavior. Consistent with this, data from Pavlovian tasks have shown that simply placing subjects into a novel context suppresses conditioned responding to associative cues, and hippocampal lesions attenuate this effect (Good et al., 1998; Penick & Solomom, 1991). In an operant/timing context, this might parralell the disruption we observed, as lesioned rats appeared less sensitive to the context-change. Future work could investigate this effect in a more targeted manner. For example, to fully dissociate the effects of context from the duration-change, one would include two further cohorts—one where the long interval changes without a context-change and another where the long-interval is held constant and a context-change occurs. This might provide a simpler assay for investigating the hippocampus’ engagement with the baseline timing network and/or how it overrides time-based behavior following context changes.

Before closing, we would like to emphasize that, while lesions only affected timing for the long cue at test, our data should not necessarily be taken to indicate that the hippocampus plays a minor or infrequent role in timing. Real-world environments are inherently unstable, particularly with compared to the tight-control experimenters enjoy in a laboratory setting. Therefore, observers must constantly adapt their expectations to changing conditions, including the context in which those changes occur. In this light, learning that a cue-duration relationship differs across two contexts might be a quite frequent and important process. Our data suggest that the hippocampus would be recruited in these cases. This highlights the importance of adapting timing tasks/designs to capture behaviors that might otherwise only occur in realistic conditions to create more ecologicallyvalid timing models.

## Supporting information

Supplemental Results

