## Supplemental Results for "The dorsal hippocampus’ role in context-based timing in rodents"

### *Unreliable CV estimates during early acquisition*

Early in the change-phase, response distributions maintained a generally early peak time, yet assumed a broad/skewed spread across the trial period (Fig S1). With respect to the Gaussian-fits, the combination of a large spread value with a small peak time and/or a general disruption to the Gaussian shape of responding resulted in implausibly large CV

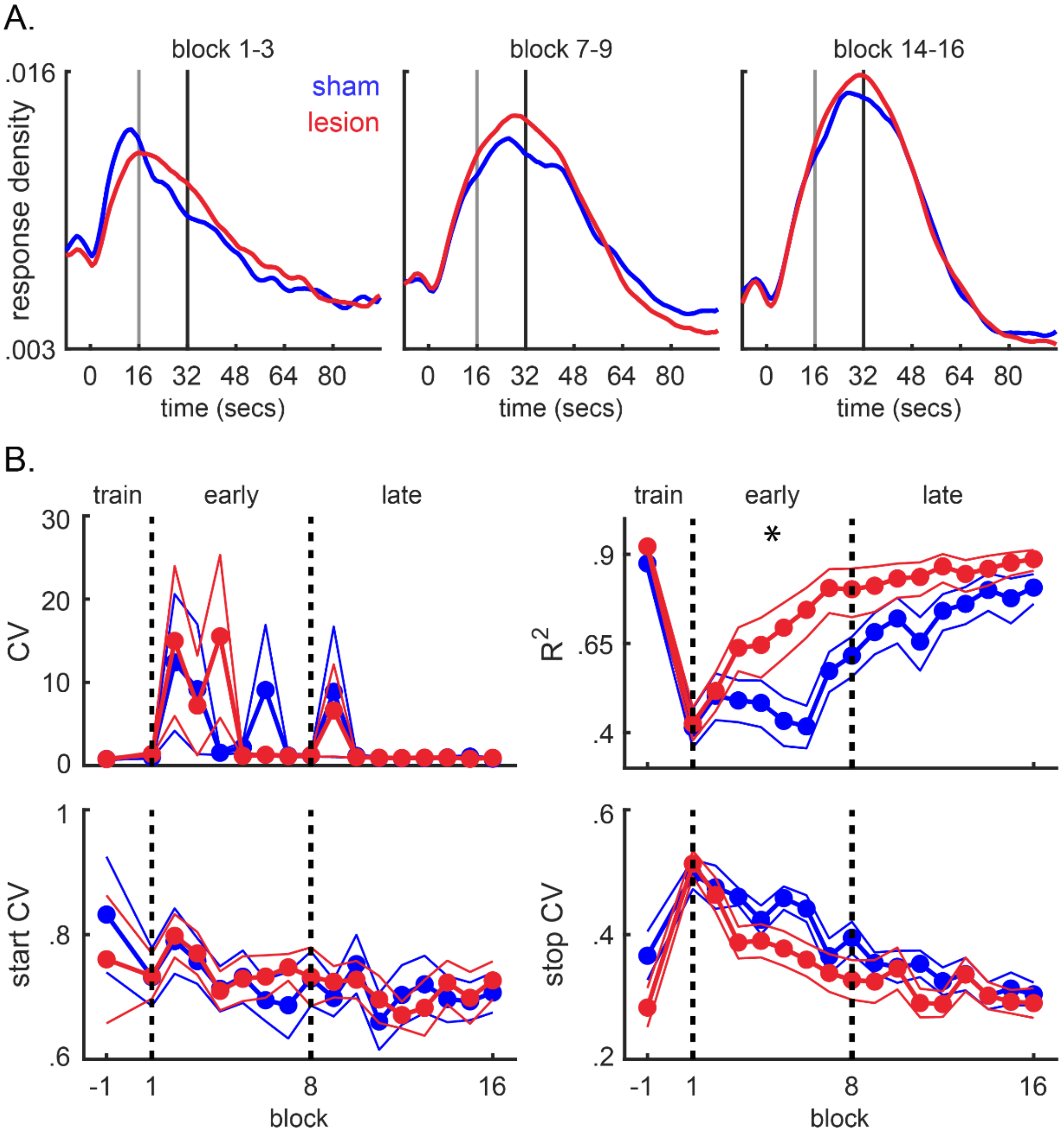


***Supplemental Figure 1.*** **Effects of dorsal hippocampus lesions on acquisition of a new interval following a context change.** Identical to Figure 3. However, the data focuses on measures of timing variability. (A) Average probe trial response rates at different points of acquisiton. Data are pooled across 3-block windows occurring at the start, middle, and at the end of the acquisiton phase (left, middle, right, respectively). Data are also smoothed over a 3-bin window, for presentation only. (B) Performance measures across individual acquisition blocks. The CV derived from the Gaussian fits is plotted at the top left and illustrates the volatility of this measure in cases where the shape of response functions was awkward spread/shape, particularly early in acquisition. For comparison, the R2 measures is still plotted to the top right, as in Figure 3. Start and stop CVs are plotted at the bottom left and right, respectively. Asterisks indicate significant group differences during a given phase.

estimates (e.g., ~10-fold increases, relative to training in some rats; Fig S1B). Therefore, we quantified the disruption in behavior with the $R^2$ of the Gaussian (Fig S1B). Furthermore, we computed CV statistics from the start and stop times. Unlike averaged response rates, this analysis can distinguish between timed and non-timed behavior trial-by-trial, rather than averaging across all responses. Consequently, these measures were more tractable, further suggesting that the source of the behavioral disruption impacting the fits did not come from time-based responding (Fig S1B). All analyses are reported in the main text.

*Single-trial analyses during context-testing*

As noted in the main text, single-trial data during testing were limited and variable. This is common during extinction in the peak-procedure, with rats showing an increase in sparse responses and/or bursts at odd times early in extinction and quickly ceasing to respond altogether (De Corte et al., 2018; Drew et al., 2017; Guilhardi & Church, 2006). When estimating the peak-time, the effect of noise is reduced by averaging across trials. However, this is obviously not possible when analyzing trials individually.

Typically, spurious start/stop times are filtered-out using a trial-exclusion criteria. A conventional approach is to only include trials where the start and stop times occur before and after the peak-time, respectively—referred to as the 'good-trials' method (Church et al., 1994; De Corte et al., 2018; Swearingen & Buhusi, 2010). This approach was generally effective in our prior report, although three rats failed to show qualifying trials in certain test conditions (De Corte et al., 2018). However, this method was too restrictive in our case, with five rats failing to meet criteria in some cases. Fortunately, laxing this criteria to trials where the start time occurred before 1.25 times the peak-time and stop times occurred after .75 times the peak time (i.e., allowing slightly later start times and earlier stop times) reduced the need for exclusion substantially (for a similar approach see Matell et al., 2016). This included one lesion rat with no qualifying trials for the short-cue in the change-context, and one sham rat for the long-cue in the original context. Therefore, we present an exploratory single-trial analysis below, only excluding those rats where needed.

In the sham group, data generally paralleled the peak-time effects, as shown in Figure S2A. Specifically, start times for both cues shifted rightward during testing across contexts [Short: Original, $t(11) = 2.54$, $p < 0.05$; Change, $t(11) = 4.5$, $p < 0.001$; Long: Original, $t(11) = 4.82$, $p < 0.001$; Change, $t(11) = 5.25$, $p < 0.001$]. Notably, these shifts were more pronounced in the change-context for both cues, indicating contextual modulation [Short: $t(11) = 2.28$, $p < 0.05$, Long: $t(11) = 3.01$, $p < 0.05$]. Moreover, relative to training, stop times shifted rightward reliably in the change-context, with no evidence of shifts during testing in the original context [Short: Original, $t(11) = 0.01$, $p = 0.999$; Change, $t(11) = 3.01$, $p < 0.05$; Long: Original, $t(11) = 0.56$, $p = 0.586$; Change, $t(11) = 7.32$, $p < 0.001$]. Consistent with a context-effect, shifts in the change-context were more pronounced than testing in the original context for the long cue [$t(11) = 9.34$, $p < 0.001$]. However, this comparison was only marginally significant for the short cue [$t(11) = 2.02$, $p = 0.069$].

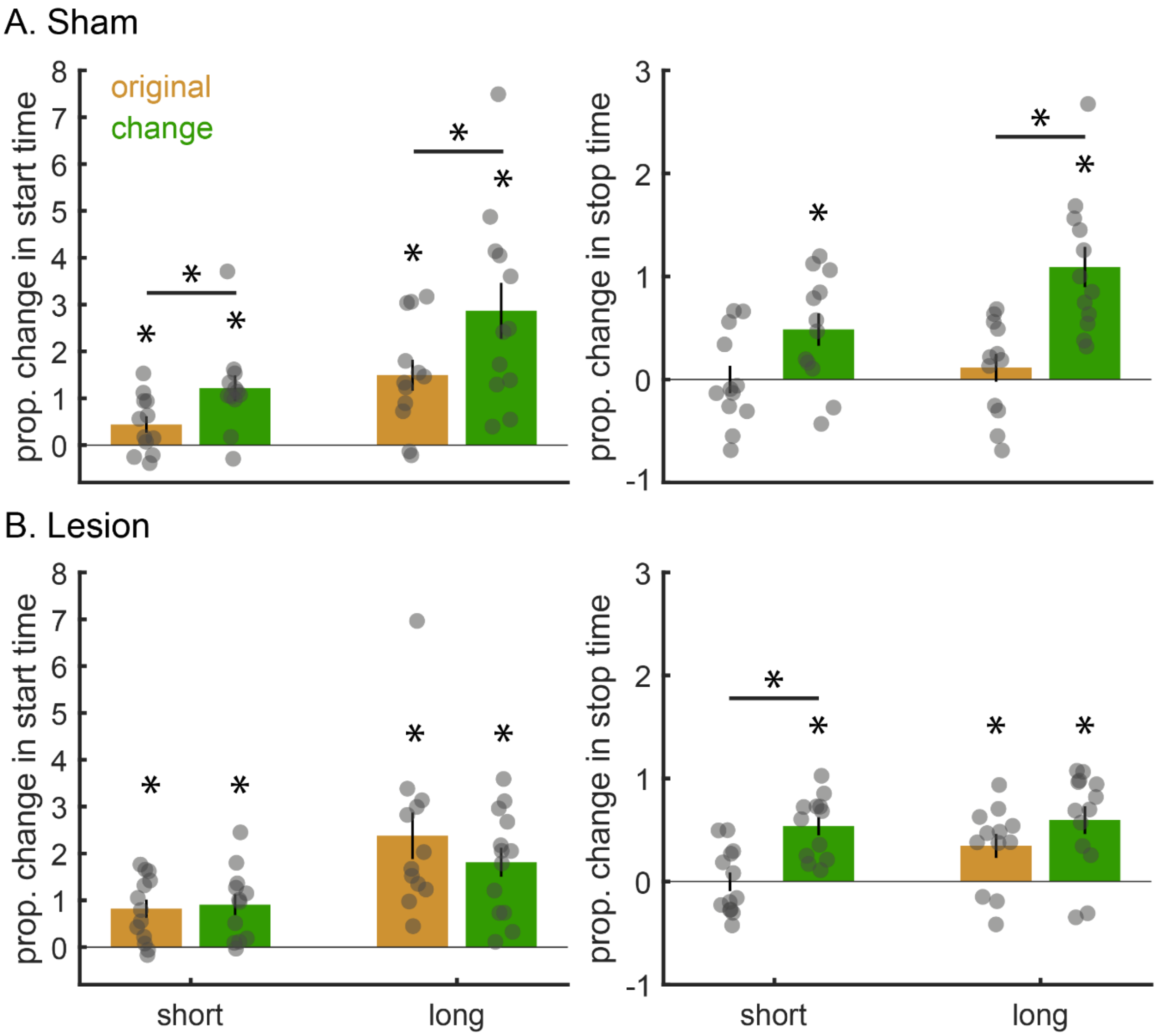


***Supplemental Figure 2. Single-trial analysis of the impact of* dorsal hippocampus lesions on context-based timing.** (A) Test data for the sham group. Proportional changes in start and stop times, relative to training, are plotted on the left and right, respectively. (B) Same format as A, yet the data now come from the lesion group. In all cases, asterisks centered over the bars indicate significant shifts from baseline training. Asterisks spanning pairs of bars indicate significant differences between testing in either context.

For the lesion group, start times also shifted rightward during testing in both contexts, relative to training [Fig S2B; Short: Original, $t(12) = 4.05$, $p < 0.005$; Change, $t(11) = 4.46$, $p < 0.005$; Long: Original, $t(11) = 6.42$, $p < 0.001$; Change, $t(12) = 6.57$, $p < 0$ .001]. Similar to peak-times, we did not observe contextual modulation for the long cue [$t(11) = -1.4$, $p = 0.19$]. However, we did not find reliable contextual modulation for short cue in this case [$t(11) = 0.12$, $p = 0.909$]. Stop times for the long cue followed the peak-time data closely, being rightward shifted during testing in both contexts with no evidence of contextual modulation [Original, $t(11) = 2.86$, $p < 0.05$; Change, $t(12) = 4.16$, $p < 0.005$; Original vs. Change: $t(11) = 0.96$, $p = 0.357$]. Importantly, for the short cue, stop times were modulated by context, with rightward shifts only occurring reliably in the change context and to a greater degree than testing in the original context [Original, $t(12) = -0.53$, $p = 0.605$; Change, $t(11) = 6.55$, $p < 0.001$; Original vs. Change: $t(11) = 4.84$, $p < 0.001$].

In relation to the peak-time results, these data provide further evidence of transfer to the short cue and contextual modulation in the sham group. In the lesion group, we continued to see transfer and a lack of contextual modulation for the long cue. However, the data may indicate that the presence or degree of contextual modulation for the short cue differs for start or stop times.
